# Vapor exposure to Δ9-tetrahydrocannabinol (THC) slows locomotion of the Maine Lobster (*Homarus americanus*)

**DOI:** 10.1101/2021.05.24.445508

**Authors:** Arnold Gutierrez, Kevin M. Creehan, Mitchell L. Turner, Rachelle N Tran, Tony M. Kerr, Jacques D. Nguyen, Michael A. Taffe

## Abstract

**Rationale:** Despite a long history of use in synaptic physiology, the lobster has been a neglected model for behavioral pharmacology. A restaurateur proposed that exposing lobster to cannabis smoke reduces anxiety and pain during the cooking process. It is unknown if lobster gill respiration in air would result in significant Δ^9^-tetrahydrocannabinol (THC) uptake and whether this would have any detectable behavioral effects.

**Objective:** The primary goal was to determine tissue THC levels in the lobster after exposure to THC vapor. Secondary goals were to determine if THC vapor altered locomotor behavior or nociception.

**Methods:** Tissue samples were collected (including muscle, brain and hemolymph) from *Homarus americanus* (N=3 per group) following 30 or 60 minutes of exposure to vapor generated by an e-cigarette device using THC (100 mg/mL in a propylene glycol vehicle). Separate experiments assessed locomotor behavior and hot water nociceptive responses following THC vapor exposure.

**Results:** THC vapor produced duration-related THC levels in all tissues examined. Locomotor activity was decreased (distance, speed, time-mobile) by 30 min inhalation of THC. Lobsters exhibit a temperature-dependent withdrawal response to immersion of tail, antennae or claws in warm water; this is novel evidence of thermal nociception for this species. THC exposure for 60 minutes had only marginal effect on nociception under the conditions assessed.

**Conclusions:** Vapor exposure of lobsters, using an e-cigarette based model, produces dose-dependent THC levels in all tissues and reduces locomotor activity. Hot water nociception was temperature dependent, but only minimal anti-nociceptive effect of THC exposure was confirmed.

## 1. Introduction

In the early fall of 2018, a minor media storm described a seafood restaurant in Maine (USA) that was proposing to expose lobsters to marijuana smoke prior to cooking (Stone, 2018). At least three testable assertions were made including 1) that some psychoactive constituent of cannabis would be transferred to the lobster via open air respiration (see follow-up reporting; (Grunewald, 2019), 2) that this would have specific behavioral effects similar to those produced in vertebrates and 3) that the cooking process would remove intoxicating psychoactive constituents from the meat thereby rendering it safe for human consumption. This latter assertion was related to a claim that “a steam as well as a heat process” would bring the lobster to 420 °F (Hinckley, 2018), which would presumably require broiling or oven baking in preference to the more typical boiling or steaming cooking method. These assertions lead to at least two key questions. Can air exposure to Δ9-tetrahydrocannabinol (THC), the primary psychoactive constituent of cannabis, produce significant tissue levels of the drug in lobsters? If so, does it have any discernible behavioral effects?

Lobsters are aquatic species that respirate via gills located inside their carapace. Lobsters can survive in air for many hours up to a few days, if they are able to keep their gills wet enough to function, but they do go into oxygen debt, e.g. across a 24 h emersion from water (Couillard and Burridge, 2015; Forgan et al., 2014). It is unclear if the gill structures would support the uptake of THC that is rendered airborne via smoke particulate or Electronic Drug Delivery System (EDDS or e-cigarette) device vapor. We recently demonstrated that vapor inhalation of THC using an e-cigarette based approach produces anti-nociceptive effects and reduces the body temperature and spontaneous activity of male and female rats (Javadi-Paydar et al., 2018; Nguyen et al., 2016). These are canonical effects that are observed after injection or oral delivery of THC in laboratory vertebrates including in rats (Taffe et al., 2015; Thompson et al., 1973), mice (Beardsley et al., 1987), monkeys (McMahon and Koek, 2007; Taffe, 2012) and dogs (Fitzgerald et al., 2013; Thompson et al., 1973), although the individual behavioral or physiological outcomes may be observed after different doses. Behavioral and physiological effects, and plasma THC levels, of THC delivered by vapor inhalation depend on the exposure duration as well as the dose administered. We have shown that dose can be controlled during inhalation exposure by varying either the concentration of THC in the e-liquid vehicle or the duration of exposure at a fixed concentration (Nguyen et al., 2016; Taffe et al., 2021). This validated platform is therefore ideal to test the hypothesis that aerosol THC exposure of lobsters has physiological effect.

Development of different animal model species, including invertebrates, for the evaluation of drug effects can offer both unique and converging advantages, as recently reviewed by Smith (Smith, 2020). The lobster is an established model for evaluating neuronal morphology, central pattern generation and synaptic mechanisms in the stomatogastric ganglion (Eisen and Marder, 1984; Marder and Eisen, 1984; Thirumalai and Marder, 2002). More practically, the lobster can be studied within institutions that are not equipped to oversee vertebrate animal research, or can be studied at reduced expense in institutions where vertebrate research is supported. A recent review indicates there are no clear data on lobster nociception and decapod crustacean investigations of nociception do not typically involve thermal stimuli (Walters, 2018); only one available report shows that crayfish are sensitive to a thermal stimulus delivered by soldering iron (Puri and Faulkes, 2015).Thus it serves the additional goal of determining if thermal nociception exists in this crustacean species which would add to the evidence for thermoception in decapod crustaceans more generally.

There is very limited evidence on whether the lobster would be behaviorally sensitive to THC exposure, however, the neuromuscular junction of lobsters appears to be regulated, in part, by cannabinoid mechanisms. Turkanis and Karler (1988) showed that THC had dose-related effects on excitatory neuromuscular junction potential amplitudes, increasing them at moderate concentrations and decreasing amplitude at higher concentrations (Turkanis and Karler, 1988). This enhances confidence that some cannabinoid-sensitive mechanisms are present in the lobster and that THC might affect locomotor behavior, despite the fact there may not be a homolog or ortholog of the vertebrate endocannabinoid receptor (CB_1_ or CB_2_) expressed in the lobster (Elphick and Egertová, 2009). The invertebrate *C. elegans* expresses a cannabinoid-like ortholog receptor (NPR-19) which mediates effects of the (mammalian) endogenous cannabinoid agonists 2-arachidonoylglycerol (2-AG) and anandamide (AEA) (Oakes et al., 2017; Pastuhov et al., 2016), suggesting that there may be as yet un-identified ortholog receptors in the lobster. It is also possible that THC acts in the lobster via transient receptor potential (TRP) channels, since THC appears to function as a ligand at TRPV2, TRPV3, TRPV4, TRPA1, and TRPV8, as reviewed (Muller et al., 2019; Starkus et al., 2019). The Caribbean spiny lobster (*Panulirus argus*) expresses 17 TRP including TRPA and TRPV homologs (Kozma et al., 2020).

Hypolocomotion is a canonical sign of cannabinoid action in rats (Tseng and Craft, 2001; Wiley et al., 2007) and mice (Wiley, 2003), and occurs after vapor inhalation of THC (Javadi-Paydar et al., 2018; Taffe et al., 2021). Thus, locomotor activity was selected to assay for evidence of *in vivo* behavioral effect. Less directly, recent studies in the crayfish, a related aquatic crustacean, have shown locomotor effects of cocaine, morphine and methamphetamine (Imeh-Nathaniel et al., 2017) and intravenous self-administration of amphetamine (Huber et al., 2018; Huber et al., 2011). This further enhances confidence that behavioral pharmacological effects of recreational drugs can be effectively assessed in the lobster.

Traditional cooking of lobster is by immersion of live animals in either boiling water or steam, leading to concerns by some that the animal might experience pain. Indeed, live cooking has been banned in Switzerland (Bachman, 2018). There is no available evidence demonstrating clearly that lobsters are sensitive to temperature, however one paper has shown that crayfish respond to a hot metal rod stimulus applied to the claw (Puri and Faulkes, 2015). Since TRPA1 homolog receptors in invertebrates are activated by high temperature, e.g., at about 48°C in one study (Caterina et al., 1997) and over 43°C in other work (Fernandes et al., 2012), these mechanisms may be the primary thermoreceptor. It is thus of interest to develop assays to determine if lobsters exhibit thermal nociceptive behavioral responses and then to determine if those responses can be altered by THC exposure. The hot-water tail withdrawal assay in rats involves a reflexive tail movement when it is inserted in hot water (∼48-52°C) and has been shown to be altered in rats after vapor inhalation of THC (Javadi-Paydar et al., 2018; Nguyen et al., 2016; Nguyen et al., 2020). Thus, one goal was to determine if warm water immersion produces a behavioral response in the lobster and if so, if THC exposure decreased thermal nociception as it does in rodents (Tseng and Craft, 2001; Wiley et al., 2007). As part of a model development, it was important to determine if different responses could be obtained from tail, claw or antenna immersion and if the response depended on the temperature of the water bath, as in (Javadi-Paydar et al., 2018).

## 2. Methods

### 2.1 Subjects

Wild caught female and male Maine lobster (*Homarus americanus;* ∼0.7-0.9 kg) were obtained from a local supermarket. When housed longer than several hours in the laboratory, the animals were maintained in chilled (∼6-10 °C), aerated aquariums (2-3 lobsters per 20 gallon tank) and fed variously with frozen krill, fish flakes and anacharis. The tissue-distribution studies were conducted under protocols approved by the Institutional Animal Care and Use Committee (IACUC) of The Scripps Research Institute due to a decision that a protocol was required for this invertebrate species. The remaining studies were conducted at the University of California, San Diego where the institution does not require protocol supervision / approval for this invertebrate species.

### 2.2 Inhalation Apparatus

Sealed exposure chambers were modified from the 259mm X 234mm X 209mm Allentown, Inc (Allentown, NJ) rat cage to regulate airflow and the delivery of vaporized drug to the chamber using e-cigarette cartridges (SMOK® TFV8 with X-baby M2 atomizer; 0.25 ohms dual coil; Shenzhen IVPS Technology Co., LTD; Shenzhen, China; controlled by e-vape controllers Model SSV-3; La Jolla Alcohol Research, Inc, La Jolla, CA, USA) as has been previously described (Javadi-Paydar et al., 2019; Javadi-Paydar et al., 2018; Nguyen et al., 2016). The controllers were triggered to deliver the scheduled series of puffs by a computerized controller designed by the equipment manufacturer (La Jolla Alcohol Research, Inc, La Jolla, CA, USA). The chamber air was vacuum controlled by a chamber exhaust valve (i.e., a “pull” system) to flow room ambient air through an intake valve at ∼1 L per minute. This also functioned to ensure that vapor entered the chamber on each device triggering event. The vapor stream was integrated with the ambient air stream once triggered. The chambers were empty of any water or bedding material for these exposures.

### 2.3 Drugs

Lobsters were exposed to vapor generated from Δ^9^-tetrahydrocannabinol (THC; 100 mg/mL) dissolved in a propylene glycol (PG) vehicle using methods previously described (Javadi-Paydar et al., 2018; Nguyen et al., 2016). One 6-second vapor puff was delivered every 5 minutes. These parameters have been developed, in previous work, to generate effective vapor fill of the chamber and significant THC-related effects in rodent subjects.

### 2.4 Tissue Collection and Analysis

For these studies, animals were obtained, dosed and euthanized for tissue collection within 4-6 hours. Lobsters were exposed to THC vapor for 30 (N=3) or 60 (N=3) minutes, then removed from the chamber and rinsed with tap water. Thereafter, they were rapidly euthanized by transection of the thoracic nerve cord using heavy kitchen shears and then transection of the thorax behind the brain by a heavy chef’s knife. Samples included the gills (N=2 for the 30-minute condition), claw muscle obtained from proximal and distal aspects, anterior and posterior segments of tail muscle, a red membrane surrounding the claw muscle (N=2 per exposure), brain, heart, liver (N=2 for the 30-minute condition) and hemolymph. Hemolymph was allowed to coagulate to facilitate analysis as ng of THC per mg of tissue, as with the other tissues. For N=2 per exposure-duration group, one claw was cooked immediately after euthanasia by boiling it in water for 10 minutes, prior to collection of muscle tissue and the red membrane that surrounds it. Tissues were frozen (−80°C) for storage until analysis was conducted. Tissue THC content was quantified using liquid chromatography/mass spectrometry (LC/MS) adapted from methods describe previously (Irimia et al., 2015; Lacroix and Saussereau, 2012). THC was extracted from brain tissue by homogenization in chloroform/ACN (Folch et al., 1957) containing 100 ng/ml of THC-d3 as internal standard (15:5:1) followed by centrifugation, decanting of the lower supernatant phase, evaporation and reconstitution in acetonitrile for analysis. Specifically, ∼200-300 mg of tissue was homogenized in 1.5 mL of chloroform, 0.500 mL of acetonitrile and 0.100 mL of deuterated internal standard (100 ng/mL THC-d3; Cerilliant). Samples were centrifuged at 3000 RPM for 10 minutes, followed by decanting of the lower supernatant phase, evaporation using a SpeedVac, and reconstitution in 200 µL of an acetonitrile/methanol/water (2:1:1) mixture. Separation was performed on an Agilent LC1100 using a gradient elution of water and methanol (both with 0.2% formic acid) at 300 µL/min on an Eclipse XDB-C18 column (3.5um, 2.1mm x 100mm). THC was quantified using an Agilent MSD6180 single quadrupole with electrospray ionization and selected ion monitoring [THC (m/z=315.2) and THC-d3 (m/z=318.2)]. Calibration curves were generated each day at a concentration range of 0-200 ng/mL with observed correlation coefficients of 0.9990.

### 2.5 Locomotion

For these studies, animals (N=7; 3 Female) were maintained in the laboratory for 4-21 days in chilled (∼8°C) aquariums. Locomotor behavior was measured in an aquatic open field arena which consisted of 45.7 cm L x 31.4 cm W x 31.4 cm D (at the bottom) clear plastic bins placed on a light surface and filled to a 20 cm depth with chilled (∼10°C) salt water and vapor exposure was to either the PG vehicle or THC on different days, with the testing order counterbalanced. The session was recorded for 30 minutes with a camera (Logitech Model #C270) placed approximately 1 m above the arena. Video recording and movement analysis was conducted with ANY-Maze (Stoelting Co.) tracking software. Parameters of movement, including total time spent mobile (seconds), total distance traveled (meters) and speed (meters/second), were extracted from the video recordings.

### 2.6 Nociception

For these studies, animals were maintained in the laboratory for 4-21 days in chilled (∼8°C) aquariums. Salt-water baths for the nociception assay were maintained at the target temperature (using placement of a beaker on a hot plate or water bath) and confirmed by thermometer immediately prior to each test. The investigator held the animal gently by the thorax and the tail or the tip of the antenna was inserted approximately 3 cm; the claws were inserted to a depth of approximately 5 cm. The latency to respond was recorded by stopwatch and a maximum 15 second interval was used as a cutoff for the assay. *Homarus* genus lobsters exhibit asymmetry of their claws with one larger (crusher) and one smaller (cutter) claw that can be on either the left or the right; feral experience appears to be necessary for proper development of the claw asymmetry as it is less pronounced in cultivated lobsters (van der Meeren and Uksnøy, 2000). This asymmetry produces a crusher muscle that is constituted of 100% slow fibers, whereas the cutter muscle exhibits only 90% fast fibers as assessed by ATPase staining and fast and slow motoneuron innervation (Govind, 1992). Thus, for this study the cutter and crusher claws were assessed independently. The order of assessment was always tail, claw, claw (cutter/crusher randomized in order) then antenna. The body part was inserted in the ambient (housing temperature, i.e., ∼-10 °C) water bath for 5-10 seconds after each warm water assessment. The temperatures for assessment were “ambient”, and then three warm temperatures (40°C, 44°C and 48°C); the order of testing of the warmer temperatures was in a counterbalanced order with at least two hours between assessments and no more than two tests per day. Lobsters were next assessed for the reaction of tail, crusher and cutter claws, and the antenna to insertion in 48°C water after vapor exposure to PG or THC for 30 or 60 minutes, with the testing order counterbalanced.

### 2.7 Data Analysis

Concentrations of THC in tissues were analyzed by mixed-effects analysis with the between-subjects factors of inhalation duration and within-subjects factor of body tissue. The nociceptive (latency) and locomotor (speed, distance, time mobile) data were assessed using within-subjects factors of vapor condition (PG vs THC), time (bin) after vapor exposure and body part (nociception). Any significant effects were followed with post-hoc analysis using Tukey correction for all multi-level, and Sidak correction for any two-level, comparisons. All analysis used Prism 9 for Windows (v. 9.1.0; GraphPad Software, Inc, San Diego CA).

## 3. Results

### 3.1 Tissue THC levels

Tissue concentrations of THC depended on the duration of vapor exposure with significantly more THC produced by 60 minutes of vapor exposure (**Figure 2**). Samples of gill were analyzed and exhibited 6,730 (SEM: 1,099) and 6,441 (SEM: 2,390) ng/g THC amounts in the 30- and 60-minute exposure conditions respectively (**Figure 2A**). This was much higher than any other tissue and is consistent with vapor containing high levels of THC collecting on the outside of the gill structure. Thus the gills were omitted from the analysis which confirmed with a main effect of exposure duration in the mixed-effects analysis [F (1, 4) = 16.97; P<0.05] of the remaining tissues (**Figure 2B**). Re-analysis with only the tissues for which N=3 were available in each exposure condition confirmed a similar main effect [F (1, 4) = 16.37; P<0.0005]. Comparison of the proximal claw THC levels with and without cooking (N=2 per exposure duration), found averages of 32.7 ng/g versus 69.5 ng/g in the 30-minute exposure and averages of 113.2 ng/g versus 478.0 ng/g in the 60-minute exposure reflecting decrements of 53% and 76%, respectively.

**Figure 1:**
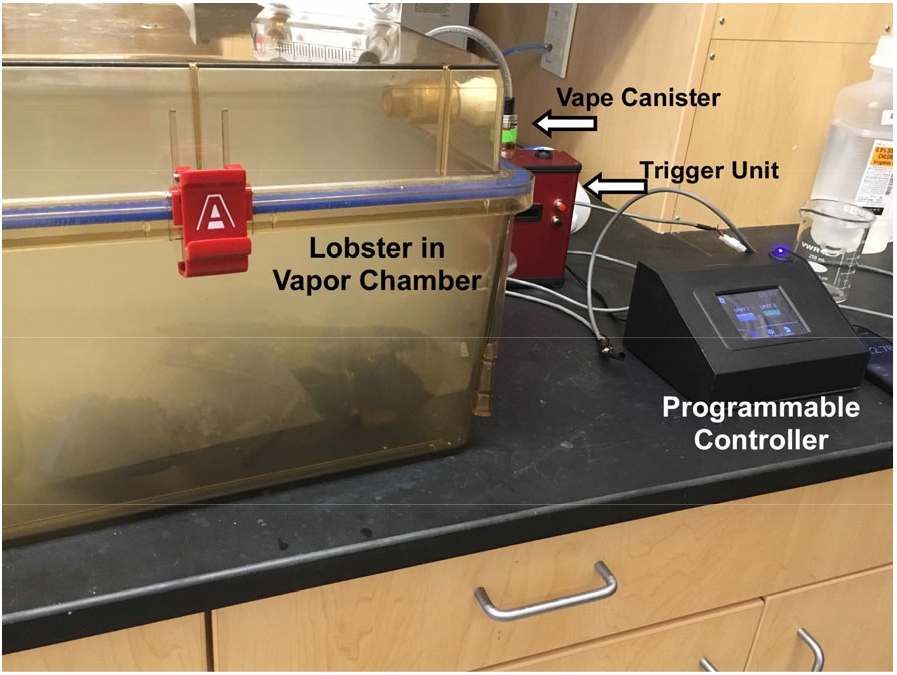
Depiction of a lobster undergoing exposure to THC vapor in a sealed chamber. Components of the vapor delivery system are identified.

**Figure 2:**
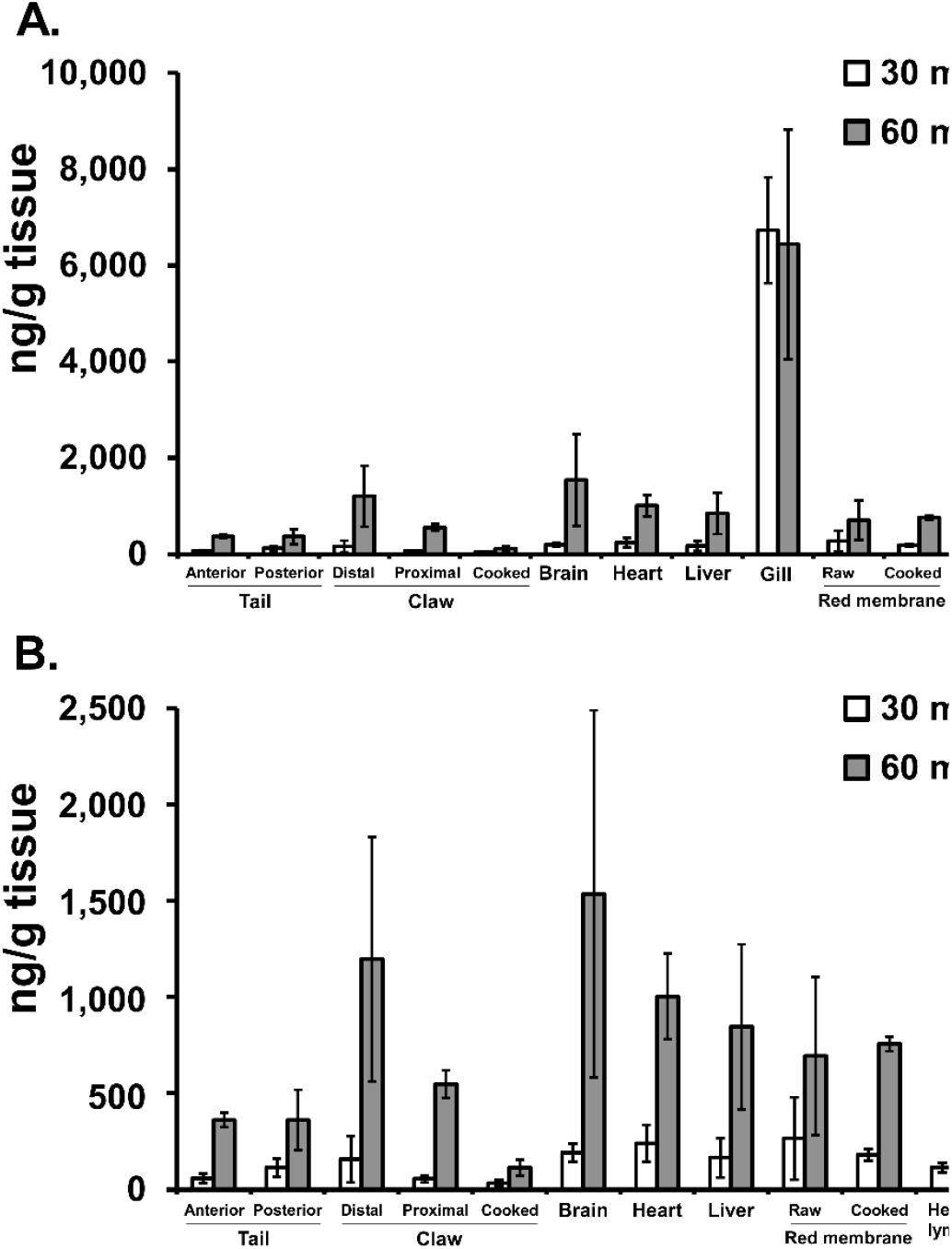
Mean (N=2-3; ±SEM) THC concentrations in various tissues after 30 or 60 minutes of exposure to THC vapor. A) all tissues and B) gill data removed to facilitate comparison of THC levels in the remaining tissues.

### 3.2 Locomotion

Lobsters spent more time mobile than immobile when in the test arena, with mean Time Mobile values in excess of 600 seconds within each 15-minute half of the session after PG exposure (**Figure 3**). Separate two-way ANOVAs for each locomotor measure confirmed a significant effect of time bin on Speed [F (1, 6) = 6.69; P<0.05], Distance [F (1, 6) = 6.74; P<0.05] and Time mobile [F (1, 6) = 15.25; P<0.01]. These analyses also confirmed a significant effect of Vapor inhalation condition on Speed [F (1, 6) = 7.83; P<0.05] and Distance [F (1, 6) = 8.68; P<0.05]. The post hoc test confirmed that Speed and Distance were lower after THC vapor exposure, compared with PG vapor exposure, for each half of the session.

**Figure 3:**
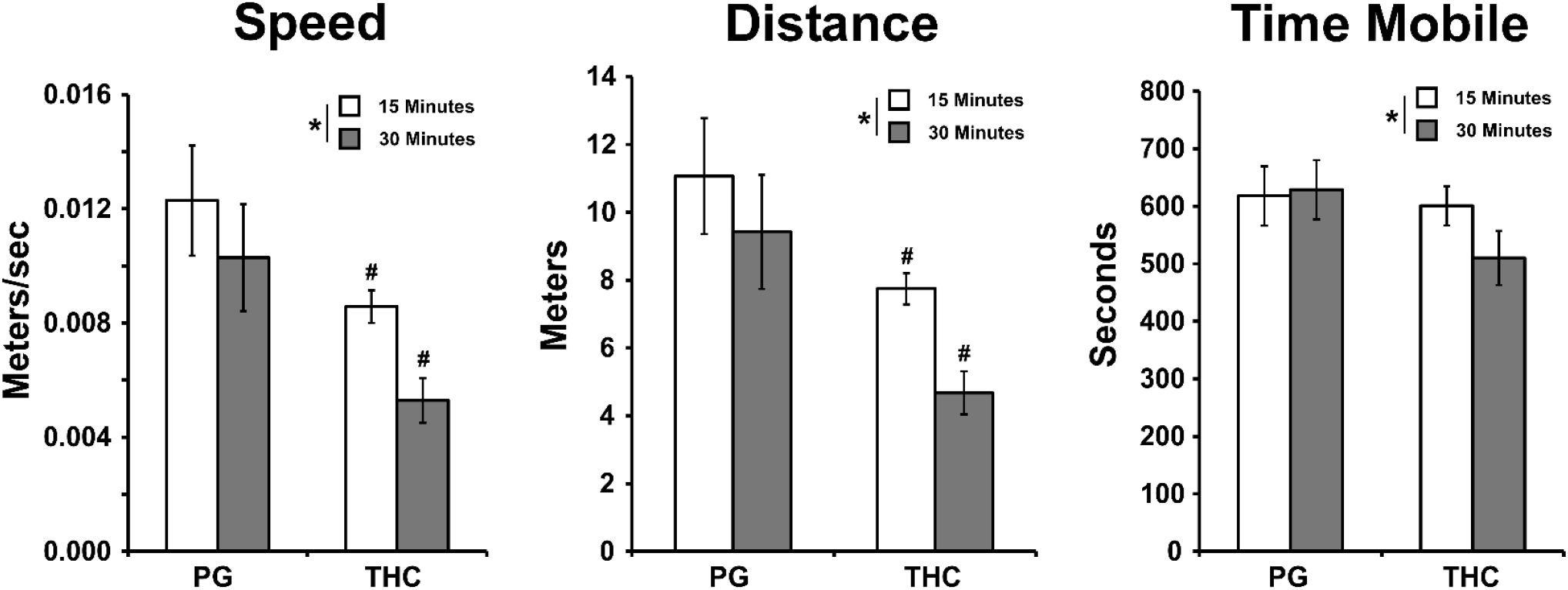
Mean (N=7; ±SEM) locomotor behavior after 30 minutes of exposure to THC vapor. Parameters of movement Speed, Distance traveled and Time in which the animal was mobile are presented. A significant difference between the first and second half of the recording interval is indicated with * and a difference relative to the PG condition, within time bin, is indicated with #.

### 3.3 Nociception

Distinct responses of tail, antenna and claw were observed following insertion in warm water but not in the ambient temperature water bath (**Figure 4**). The response following insertion of the tail consisted of two distinguishable responses. Sometimes, a reflexive and powerful contraction of the tail muscle was observed first (see **Supplemental Materials**). This appeared to be the caridoid escape response best described in crayfish, but likely present in the lobster (Lang et al., 1977; Otsuka et al., 1967), which is a complex behavior mediated by lateral giant and medial giant interneurons (Wiersma and Ikeda, 1964). In other cases, the lobster initiated distinct movements of legs and claws, this often preceded the powerful tail contraction. Thus, the tail assay was scored with two latency values, the very first reaction of any type (tail reaction) and the tail contraction if it occurred. For analysis, the time of first overt response (“tail reaction”) was used. The ANOVA confirmed a significant effect of body part [F (2.495, 22.45) = 11.94; P<0.0005] and water temperature [F (2.668, 24.01) = 30.89; P<0.0001] on withdrawal latency. The post hoc test of the marginal means confirmed that withdrawal latencies in ambient temperature differed from all other temperatures, and latencies in each of 44°C and 48°C differed from latencies observed after 40°C insertion. Post hoc of the marginal means for body part confirmed latency for the crusher claw was slower than for each of the other body parts.

**Figure 4:**
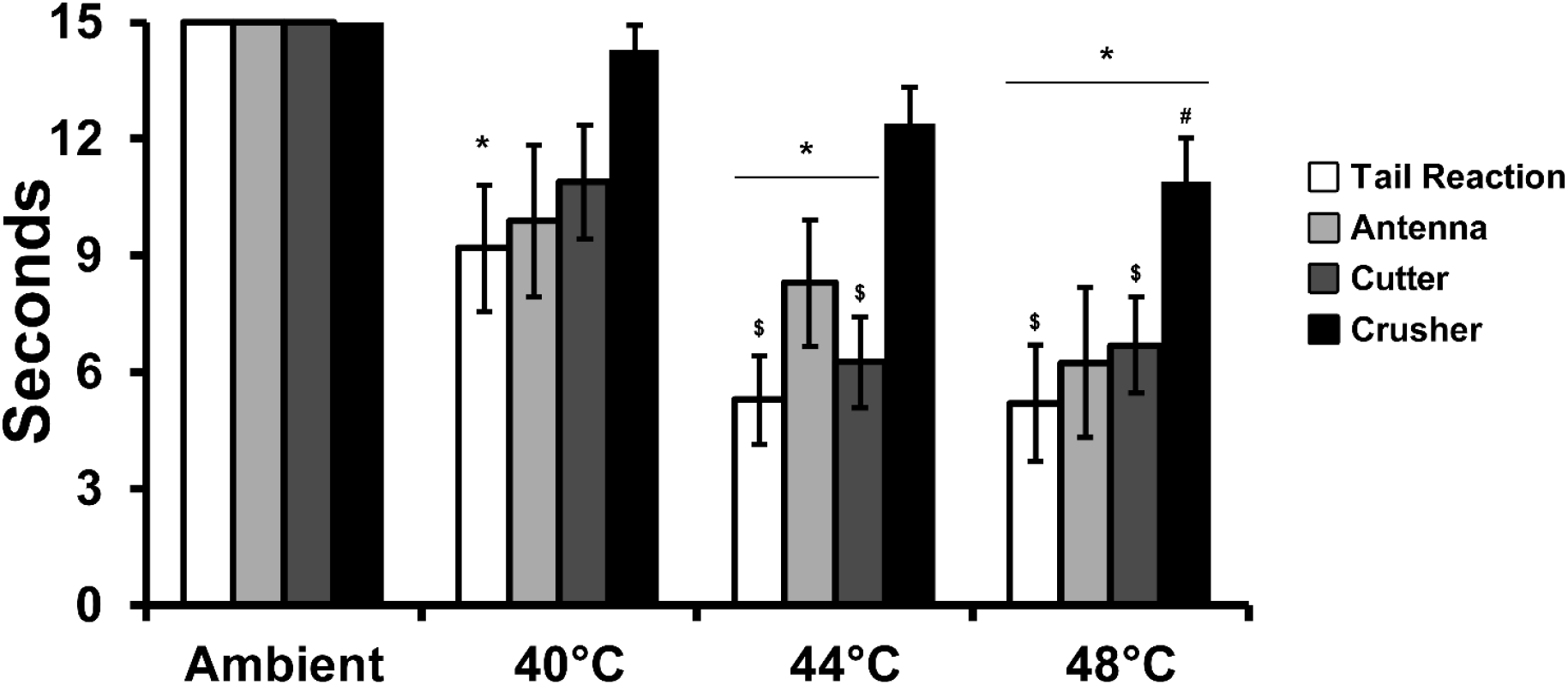
Mean N=11 (3 Female); latency to respond to the immersion in the water bath of the indicated temperature. A significant difference from ambient, within body part, is indicated with *, a difference from the 40°C condition, within body part with # and a difference from the crusher claw, at a given temperature, with $.

The reaction to the 48°C water insertion was only marginally affected by vapor exposure to THC (**Figure 5**). The three-way ANOVA confirmed a significant effect of body part [F (3, 24) = 4.59; P<0.05], and of the interaction of Vapor Condition with Exposure Duration [F (1, 24) = 6.68; P<0.05] on reaction latency. Follow-up two-way ANOVAs for each exposure duration confirmed a significant effect of Body Part [F (1.937, 11.62) = 8.03; P<0.01] after 30 minutes of exposure (but no effect of Vapor Condition) and a significant effect of Body Part [F (1.761, 10.57) = 6.28; P<0.05] and the interaction of Body Part with Vapor Condition [F (2.477, 14.86) = 4.19; P<0.05] after 60 minutes of exposure. The post-hoc test failed to confirm any significant difference associated with Vapor Condition for any individual body part.

**Figure 5.**
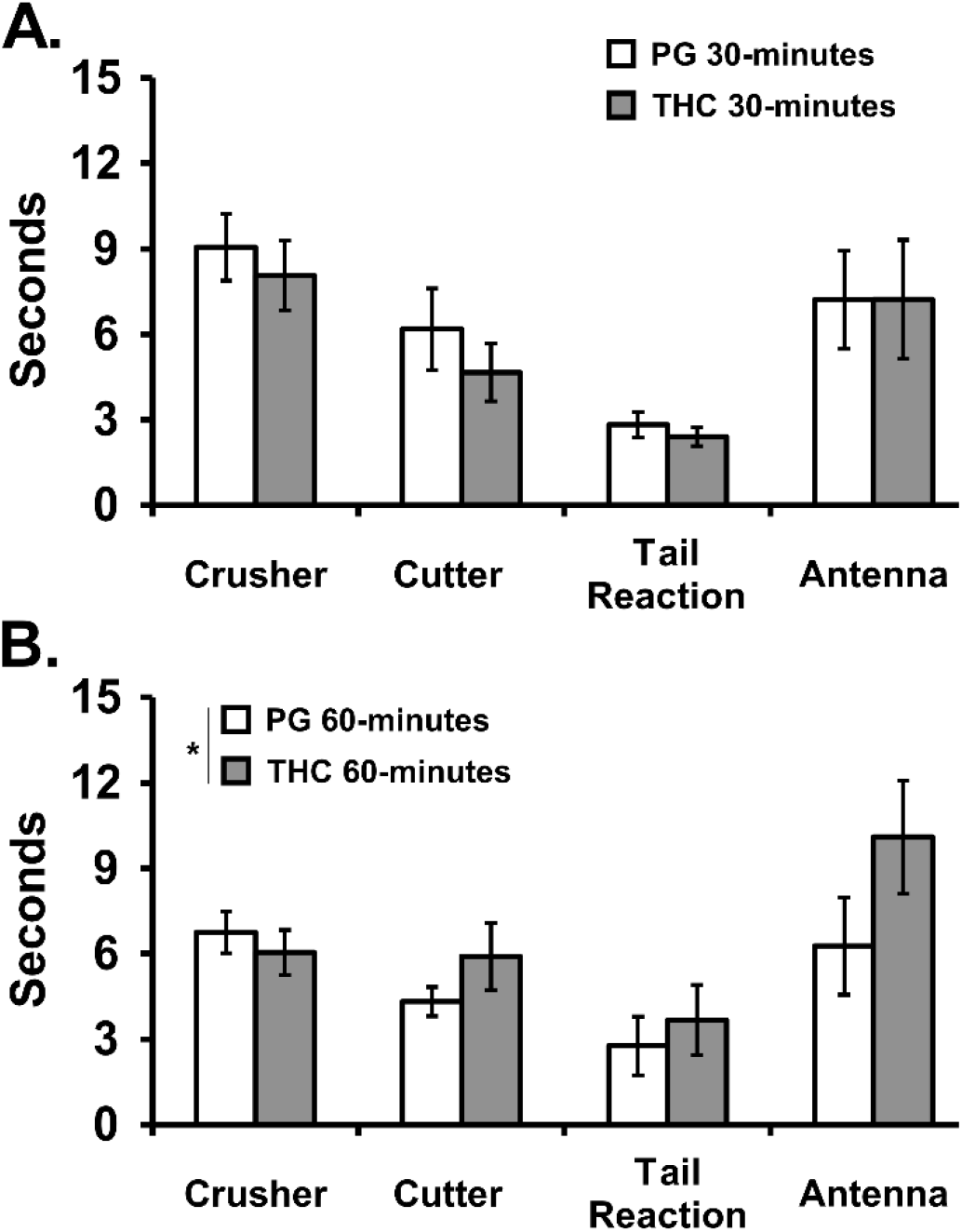
Mean (N=7; ±SEM) latency to react to immersion in 48°C water after A) 30 or B) 60 minutes of exposure to vapor from the propylene glycol (PG) vehicle or THC (100 mg/mL in the PG). Tail rxn = Tail reaction, i.e., the first defined movement. A significant difference between Vapor Conditions, across body part, is indicated with *.

## 4. Discussion

The primary finding of this study was that vapor exposure of Maine lobsters (*Homarus americanus)* to Δ9-tetrahydrocannabinol (THC), using an e-cigarette based system, produces tissue levels of THC in a dose (time of vapor exposure) dependent manner. THC was confirmed in the hemolymph (the “blood” of the lobster), claw and tail muscle, brain, heart and liver (**Figure 2**). This wide distribution across body tissues is consistent with respiratory uptake, i.e., via the gills with distribution by the hemolymph circulation of the lobster. This conclusion is further supported by the much higher amount of THC that was associated with the gill tissue, consistent with a limited uptake by the respiration system of the lobster.

The THC exposure also had behavioral consequences, since locomotor activity was significantly reduced after exposure to THC vapor compared with exposure to the vehicle vapor (**Figure 3**). Hypolocomotion is a canonical feature of THC exposure in rats and mice, at least at higher doses, thereby confirming a similarity of effect across the vertebrate and invertebrate organisms in which this has been evaluated. Thus, the assertions of the restaurateur that cannabinoids could be introduced into the lobster by atmospheric exposure (Grunewald, 2019; Hinckley, 2018; Stone, 2018), and that this would be in sufficient amount to induce behavioral effect, is supported. The impact of THC on thermal nociception was, however, minimal.

The locomotor effects may be specifically related to a report of THC altering the amplitude of excitatory potentials at the lobster neuromuscular junction in a concentration dependent manner (Turkanis and Karler, 1988). Overall, however, it confirms that the levels of THC achieved by as little as 30 minutes of vapor exposure were behaviorally significant. One caveat for the locomotor studies is that the arena was not as large compared with the size of a lobster as the similar ratio for typical open field studies conducted in rodents. Similarly, the water depth was limited to that necessary to cover the lobster to facilitate the video tracking for this initial investigation. Nevertheless, the animals were able to express movement, turn around, change direction, etc., and traveled about 20 meters after the vehicle exposure condition. It would be of interest in future studies to assess locomotor behavior in a larger arena or to assess behavior in a deeper aquatic environment.

In the nociception experiment, lobsters were observed to respond to warm water immersion of claw, tail or antenna in a temperature-dependent manner (**Figure 4**). This provides evidence of thermal nociception in the lobster for the first time (Walters, 2018), and is consistent with prior work which has shown thermal nociception in crayfish, using a warm (54°C) metal stimulus on the claw and antenna (Puri and Faulkes, 2015). No response to immersion in maintenance temperature (∼10°C) salt water was observed in this study, using the 15 sec cutoff. (In initial/pilot studies, there was also no response observed at laboratory ambient temperature of ∼22-24°C). At temperatures from 40-48°C, however, the lobsters made distinct motor responses upon immersion of the tail, the claws or the antenna. Tail immersion resulted in a clear response of legs and claws and/or a strong flick of the tail which was in many cases repetitious (see **Supplementary Materials**). These latter behaviors can be considered within escape responses of lobsters and crayfish, with evidence from the latter more plentiful. In the crayfish, sudden onset stimuli evoke tail movements associated with lateral and/or medial giant neuronal activity whereas more gradual stimuli evoke tail movements which do not involve the lateral or medial giant (Wine and Krasne, 1972). This study observed a range of apparent behavioral responses including gradual movements, strong flips and repeated flipping of the tail suggesting a diversity of sensory experiences from the sudden to the gradual. Additional experimentation would be required to further dissociate the thermoceptive responses under various conditions but the critical factor for this study was the detection of the hot water stimulus. Immersion of the claws or antenna also resulted in a distinct movement to remove the appendage from the water, consistent with thermoception. Temperature dependent differences in response latency were observed for the warm water challenges, with 40°C less noxious than 44 or 48°C. This graded response is what is observed with a similar nociceptive assay in rats (Javadi-Paydar et al., 2018) and further enhances confidence in the specificity of the response to the noxious stimulus. The pronounced difference in sensitivity between the crusher and cutter claws provides another important validation of the model. Prior reports have focused on claw morphology and muscle fiber type (van der Meeren and Uksnøy, 2000) and this extends this by demonstrating a clear behavioral insensitivity of the crusher compared with the cutter claw. Finally, the effect of THC vapor exposure on thermal nociception was minimal under the tested conditions. Surprisingly, despite the locomotor effect of 30 min of THC vapor exposure, there was no impact relative to vehicle vapor exposure on the latency of the response to warm water immersion (of any body part). It required 60 minutes of exposure to THC to produce any significant effect (**Figure 5B**), which was very small in magnitude. Although THC has limited anti-nociceptive impact in rodents relative to opioids (Gutierrez et al., 2021; Nguyen et al., 2019) and has a limited dose-effect range due to this low ceiling, it is typically more robust in rodents than what was observed here.

The present results and the work on lobster neuromuscular junctions (Turkanis and Karler, 1988) suggest that THC has specific neuropharmacological effect, however, there does not appear to be a vertebrate endocannabinoid receptor (CB_1_ or CB_2_) expressed in the lobster or crayfish (Elphick and Egertová, 2009); this is similar to behavioral effects in *Drosophila melanogaster* (He et al., 2021) which likewise lack CB_1/2_ analogs. This turns attention to the possibility that THC acts in the lobster via temperature activating (Chen, 2015) transient receptor potential (TRP) channels. THC appears to function as a ligand at TRPV2, TRPV3, TRPV4, TRPA1, and TRPV8, as reviewed (Muller et al., 2019; Starkus et al., 2019), and the Caribbean spiny lobster (*Panulirus argus*) expresses 17 TRP channels including TRPA and TRPV homologs (Kozma et al., 2020). A further clue is provided by the fact that another cannabis constituent which does not have substantial activity at endocannabinoid receptors (Boggs et al., 2018), cannabidiol (CBD), also had inhibitory effects in Turkanis and Karler (1988), and CBD modulates several THC sensitive TRPs (Muller and Reggio, 2020; Starkus et al., 2019). One apparent concern for this interpretation is that mammal TRPA1 is activated by noxious low, but not high, temperature. Nevertheless, thermosensitive TRPs may be a particularly good candidates for any anti-nociceptive effects in decapod crustaceans given that TRPA1 from invertebrates (and some non-mammalian vertebrates) are indeed activated by high temperature. In some work, TRPV1 activates at about 48 °C (Caterina et al., 1997), in other work over 43 °C as reviewed (Fernandes et al., 2012). There may be species differences in the activation temperature and as a comprehensive review observes that while TRPA1 activation temperatures vary across species/ortholog, activation is always above the preferred temperature of the species (Laursen et al., 2015). Overall, the likely activation temperature for TRP homologs in the lobster are likely to accord with the temperature range established here for thermal nociception, i.e., observed at 40°C and above, but not at 22°C (**Figure 4**).

A prior finding, however, that capsaicin did not influence the apparent thermal nociceptors in crayfish (Puri and Faulkes, 2015) may suggest that whatever small effect was produced on nociception here was not mediated by a capsaicin sensitive TRP channel, such as TRPA1 or TRPV1. Alternately, the fact that TRPV1 can desensitize to capsaicin (Fernandes et al., 2012) may have resulted in an apparent divergent result for noxious heat and capsaicin stimuli, presumably as a consequence of the specific methods used and potentially the species/ortholog in question. Finally, it is possible that there may be an as yet undiscovered ortholog of mammalian cannabinoid receptors in the lobster. For example, the invertebrate *C. elegans* expresses a cannabinoid-like receptor (NPR-19) that appears to mediate effects of the endogenous cannabinoid agonists 2-arachidonoylglycerol (2-AG) and anandamide (AEA) (Oakes et al., 2017; Pastuhov et al., 2016).

There are a few limitations to this study that may be useful to address in any future work. In the lobster behavior experiments, housing for the range of 4-21 days prior to a given assessment was never explicitly tested for potential effects. No major changes were noticed, but it is not impossible this would contribute. Likewise, we selected a fixed housing temperature, within the range described for this species in their natural habitat, so any effect of housing at one or the other end of their temperature range was not determined. As mentioned above, the locomotor assessment was conducted in an arena too small to divide into zones, e.g. Center vs Periphery, that in the context of a rodent test would permit assessment of anxiety-like avoidance of the center; a larger arena might facilitate such investigations. The nociception assay provides strong evidence for detection of a warm water stimulus, however it is possible that additional analysis of the response of the tail would provide further insight. Because the main goal was to determine if any response would be made, and there is no available information on how lobsters would respond to a thermally noxious stimulus, it was decided to operationalize the first clearly detectable response as the target latency. Future studies which determine the consistency of the strong flip, repeated flipping (see Supplementary Materials) or the slower withdrawal response, both between and within individuals, would further define this response.

In conclusion, these data confirm a method for studying the effects of aerosol THC exposure in a lobster model. Duration-dependent levels of THC were observed in the species’ tissues and a reduction in locomotor behavior was produced. The animals also responded in a temperature-dependent manner to the immersion of tail, claw or antenna in a hot water bath, indicating thermal nociception. This latter conclusion was further enhanced by the observation of differential sensitivity in the cutter and crusher claws. Further experimentation will be required to fully investigate other behavioral outcomes, including anxiety-like measures.

## Acknowledgements

These studies were supported in part by USPHS grants R01 DA035281; R01 DA035482 and R44 DA041967; all animals were purchased with non-NIH funds. The NIH/NIDA had no role in study design, collection, analysis and interpretation of data, in the writing of the report, or in the decision to submit the paper for publication. La Jolla Alcohol Research, Inc (LJARI) engages in commercial development of vapor inhalation techniques and equipment, including with support from the R44 DA041967 SBIR grant. LJARI donated funds for some of the lobster purchases but was not directly involved in the design of the experiments, analysis and interpretation of data or the decision to submit the study for publication. The authors declare no additional financial conflicts which affected the conduct of this work. The authors are grateful to Zen Faulkes, Ph.D., Professor in the School of Interdisciplinary Science, McMaster University, for advice on the handling and euthanasia of crustacean species. The authors are likewise grateful for comments made on Twitter in response to the initial pre-print posting (https://www.biorxiv.org/content/10.1101/2021.05.24.445508v1) of this work (including contributions from @AizenmanLab, @boehninglab, @DoctorZen, @drosophilosophy, @Hoosierflyman, @StolfiAlberto, and @millerlab, among others) and by peer reviewers for substantially improving the scholarship.

## Author Contributions

AG contributed to the overall design of the studies, to assay development, created the locomotor assessment approaches and video analysis, collected data, and provided initial data analysis. KMC contributed to behavioral assay development and study design, euthanized lobsters and collected tissues, designed and implemented housing and husbandry procedures, collected data and performed initial data analysis. MLT and RNT made critical contributions during assay development, including the critical cutter/crusher claw distinction, collected data and assisted with assessing relevant literature. TMK conducted tissue THC assays. JDN assisted with vapor exposure apparatus, overall experimental design and data collection. MAT secured the animal protocol approvals, euthanized lobsters, collected tissues, designed and conducted the cooking assay, contributed to the overall design of studies, conducted statistical analysis, figure creation and initial drafting of the manuscript. All authors reviewed drafts and approved the manuscript.

